# Systematic Optimization of Activity-Based Protein Profiling for Identification of Polysorbate-Degradative Enzymes in Biotherapeutic Drug Substance Down to 10 ppb

**DOI:** 10.1101/2024.08.29.610306

**Authors:** Taku Tsukidate, Anita P. Liu, Shannon Rivera, Alyssa Q. Stiving, Jonathan Welch, Xuanwen Li

**Author notes:** **Corresponding Author** XL.

## Abstract

The identification and control of high-risk host cell proteins (HCPs) in biotherapeutics development is crucial for ensuring product quality and shelf life. Specifically, HCPs with hydrolase activity can cause degradation of excipient polysorbates (PS), leading to a decrease in the shelf life of the drug product. In this study, we systematically optimized every step of an activity-based protein profiling (ABPP) workflow to identify trace amounts of active polysorbate-degradative enzymes (PSDEs) in biotherapeutics process intermediates. Evaluation of various parameters during sample preparation pinpointed optimal pH level and fluorophosphonate (FP)-biotin concentration. Moreover, the combined use of a short liquid chromatography gradient and the fast-scanning Parallel Accumulation Serial Fragmentation (PASEF) methodology increased sample throughput without compromising identification coverage. Tuning Trapped Ion Mobility Spectrometry (TIMS) parameters further enhanced sensitivity. In addition, we evaluated various data acquisition modes including PASEF combined with data-dependent acquisition (DDA PASEF), data-independent acquisition (diaPASEF), or parallel reaction monitoring (prm-PASEF). By employing the newly optimized ABPP workflow, we successfully identified PSDEs at a concentration as low as 10 parts per billion (ppb) in a drug substance sample. Finally, the new workflow enabled us to detect a PSDE that could not be detected with the original workflow during a PS degradation root-cause investigation.

## Introduction

Biotherapeutics, such as therapeutic monoclonal antibodies (mAbs) products, have become a predominant therapeutic modality for the treatment of a wide range of diseases. The global sales revenue for all mAb drugs alone totaled nearly $115.2 billion in 2018 and is expected to reach $300 billion by 2025.^1,2^ Biotherapeutics produced in host cells, such as Chinese hamster ovary (CHO), go through downstream purifications to remove product and process related impurities, such as host cell proteins (HCPs). The identification and control of those high-risk HCPs is critical to biotherapeutics development.^3^ For example, certain HCPs with hydrolase activity can cause formulation excipient polysorbate degradation if they are not effectively removed during downstream purification, which can significantly shorten the shelf life of the drug product.^4^ If proven in a manufacturing setting even a modest 5–10 % increase in product shelf life resulting from the reduction and control of HCPs would translate to millions of dollars of cost saving, not to mention significant improvement in downstream purification process specifically designed to remove those high-risk HCPs. The increased product shelf life may hopefully reduce the drug cost and improve the equitable access to those life-saving medicines.

Liquid chromatography tandem mass spectrometry (LC– MS/MS)–based proteomics has proven to be a powerful approach to HCP analysis which enables tracking of individual HCPs throughout a purification process.^5,6^ HCP analysis poses a unique challenge to conventional abundance-based proteomics because of the large dynamic range of samples that can span more than six orders of abundance between the therapeutic antibody and low-abundance HCPs. Accordingly, the past several years saw the development of various sample preparation methods such as native digestion^7,8^, molecular weight cutoff^9^, and ProteoMiner enrichment^10^ that improved sensitivity for low-abundance HCPs in the range of 0.05–1 ppm. However, abundance-based proteomics does not offer information about the functional state of HCPs that would help determine the cause of an issue at hand such as excipient degradation. In contrast, activity-based protein profiling (ABPP) employs active site–directed chemical reporters to selectively enrich active enzymes in complex samples.^11,12^ Thus, we and others have previously reported ABPP protocols for HCP analysis which enables sensitive detection and functional assessment of specific HCPs.^13,14^ More recently, Liu et al. developed a polysorbate-inspired chemical reporter and demonstrated the detection of polysorbate-degradative enzymes (PSDEs) down to 80 ppb.^15^ Herein, we describe further optimization of the ABPP approach and demonstrate confident identification of active PSDEs at the 10-ppb level (**Figure 1**). The established ABBP workflow will fill the gap between PSDE abundance and activity, allowing more meaningful polysorbate degradation investigations for biotherapeutic development.

**Figure 1.**
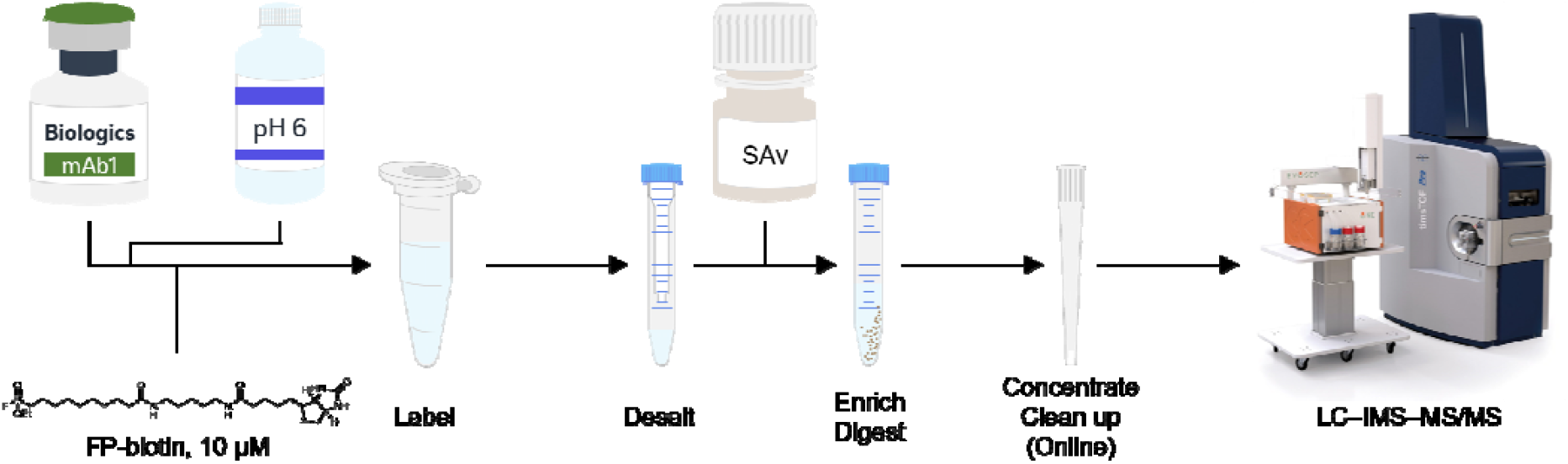
Overview of ABPP of biotherapeutics samples with FP-biotin. Materials and methods

### Protein samples

Recombinant PLA2G7 and SIAE proteins were custom ordered from GenScript. mAb1 harvested cell culture fluid (HCCF), mAb1 Protein A pool (PAP), mAb1 drug substance (DS), and mAb2 ultrafiltration pool (UFP) were internally sourced.

### ABPP sample preparation from HCCF

HCCF (100 μL, ∼ 0.5 mg) was buffer-exchanged with either 100 mM sodium acetate pH 5, 100 mM histidine pH 6, DPBS pH 7, or 50 mM tris-HCl pH 8 through a Zeba Spin Desalting Column, 7K MWCO, 0.5 mL (Thermo Fisher Scientific) according to the manufacturer’s instructions. The buffer-exchanged material was incubated with fluorophosphonate (FP)-biotin (WuXi AppTec; 0, 0.1, 1, or 10 μM) at 25 °C for 0.5 or 2 h and then buffer-exchanged with DPBS pH 7 through a Zeba Spin Desalting Column to remove excess FP-biotin and adjust pH. To this were added 0.2 % SDS / DPBS (900 μL) and Pierce Streptavidin Magnetic Beads (20 or 40 μL; Thermo Fisher Scientific). After 1-h incubation at room temperature, the beads were washed three times with 0.2 % SDS / DPBS pH 7 (1 mL) and three times with 1 M urea / water (1 mL). Then the beads were incubated with 0 or 10 mM TCEP-HCl or 0 or 20 mM iodoacetamide at 25 °C for 0.5 h and washed twice with 50 mM tris-HCl pH 8 (1 mL). The beads were incubated with 50 mM tris-HCl pH 8 (50 μL) containing 0.1 μg trypsin (Promega) at 25 °C for 20 h. The supernatant (∼ 50 μL) was cleaned up with Pierce C18 Spin Column (Thermo Fisher Scientific) according to the manufacturer’s instructions, vacuum-dried, and reconstituted with 0.1 % FA / water (5 μL). Two μL was injected. Alternatively, in our optimal protocol, HCCF (100 μL) was directly incubated with FP-biotin (10 μM) at 25 °C for 1 h and filtered through a Zeba Spin Desalting Columns. The filtrate was diluted with 0.2 % SDS / DPBS (900 μL) and incubated with Pierce Streptavidin Magnetic Beads (20 μL) at room temperature for 1 h. The beads were washed three times with 0.2 % SDS / DPBS pH 7 (1 mL) and three times with 1 M urea / water (1 mL), followed by incubation with 50 mM tris-HCl pH 8 (50 μL) containing 0.1 μg trypsin (Promega) at 25 °C for 20 h. The supernatant (20 μL) was loaded on Evotip Pure (Evosep) according to the manufacturer’s instructions.

### ABPP sample preparation from PAP, UFP or DS

PAP, UFP or DS (5 mg) was diluted to 5 g/L with 100 mM histidine pH 6 and was incubated with 10 μM FP-biotin at 25 °C for 1 h and filtered through a Zeba Spin Desalting Column, 7K MWCO, 2 mL (Thermo Fisher Scientific). To this were added DPBS (4 mL), 10 % SDS (100 μL), and streptavidin magnetic beads (200 μL). After 1-h incubation at room temperature, the beads were washed three times with 0.2 % SDS / D**PB**S (5 mL) and three times with 1 M urea / water (5 mL). T**hen** the beads were transferred to a microcentrifuge tube using **50** mM tris-HCl (1 mL) and incubated with 50 mM tris-HCl pH **8** (50 μL) containing 1 μg trypsin at 25 °C for 20 h. The supe**rna**tant (∼ 50 μL) was loaded on Evotip Pure (Evosep). Alterna**tiv**ely, mAb2 UFP (20 mg) was processed as we previously de**scr**ibed for comparison.^13^

### LC–MS analysis

Peptides were separated on the Evosep One LC system^16^ with Pepsep C18 columns (Bruker) using the manufacturer’s pre-defined methods denoted by daily sample throughput e.g., 30 samples per day (SPD) and analysed on the Bruker timsTOF Pro 2 system. The DDA PASEF method was optimized based on the standard DDA PASEF method (“DDA PASEF-standard_1.1sec_cycletime.m”). The diaPASEF method was optimized as we previously described and covered 300–1200 m/z with 25 diaPASEF scans and two ion mobility windows per diaPASEF scan (i.e., 50 variable windows).^6^ The prm-PASEF method targeted all precursors that were identified for PLA2G7 and SIAE in the DDA PASEF analysis of the mAb1 HCCF ABPP sample. The time and mobility scheduled acquisition boxes were set with 60 s of tolerance on retention time and 0.05 1/*K*_0_ on ion mobility. In all methods, the ion mobility range was set as 0.7–1.43 Vs cm^−2^ with 100 ms of accumulation time and 100 ms of ramp time unless otherwise noted. Alternatively, peptides were separated on the EASY-nLC 1200 system and analyzed on Q Exactive HF-X system as we previously described.^13^

### Data analysis

DDA PASEF data were processed with MaxQuant v2.4.10 or v2.6.1^17^. diaPASEF data were processed with DIA-NN v1.8.1 or v1.9.1^18^. Protein quantification was performed in R version 4.0.3 using the fast_MaxLFQ function from the iq package^19^ based on the Precursor.Normalised column of DIA-NN main reports, which represents the aggregated and normalized MS2 quantities. Prm-PASEF data were processed with Skyline v24.1.0.199^20^. Precursors were filtered by requiring isotope dot product > 0.9 and their intensities were calculated by summing all associated fragment peak areas and subtracting background signals. Protein intensities were then calculated as sums of the intensities of the three most abundant precursors. The precursor and protein FDR thresholds were set at 1 % for HCCF- or PAP-based samples and at 5 % for UFP- or DS-based samples.^21^ The FrF2^22^ and rsm^23^ packages were used to generate and analyze the Placket-Burman screening design and central composite design, respectively.

### High-performance liquid chromatography–charged aerosol detection (HPLC-CAD) for PS degradation analysis

HPLC-CAD was performed on Waters 1200 Series and Thermo Corona Charged Aerosol Detector with Waters Oasis Max column and a 20-min gradient. Mobile phases were (A) 0.5 % acetic acid in water and (B) 0.5 % acetic acid in isopropanol.^24^

## Results and discussion

The active site–directed chemical reporter FP-biotin targets over 100 murine and human metabolic serine hydrolases (mSHs) in cell and tissue samples.^25,26^ While the standard protocol labels mSHs with FP-biotin at pH 7–8^27^, the optimal pH level for enzymatic activity varies for different mSHs and can affect their FP-biotin labeling levels^28,29^. Other sample preparation parameters such as FP-biotin labeling time, bead volume, reductant and alkylation reagent concentrations could also impact assay sensitivity as well as operational cost and/or hands-on time. To identify sample preparation parameters with significant effect on sensitivity towards various mSHs, we performed a twenty-sample Plackett–Burman screening experiment^30^ including six aforementioned parameters at two levels (**Table 1**). Indeed, varying pH level resulted in differential enrichment of several mSHs (**Figure 2A**). For example, lysosomal mSHs, namely PRCP and PLA2G15, were significantly enriched in samples labelled at pH 5 whereas several other mSHs including LPL, IAH1 and SIAE were significantly enriched in samples labelled at pH 8, which is consistent with their optimal pH levels for activity^29,31^. In addition, the higher FP-biotin concentration (10 μM) increased protein intensities selectively for mSHs compared to using the lower FP-biotin concentration (1 μM), indicating limited off-target labelling of other inactive proteins. On the other hand, the other four parameters had negligible effect. These results suggest that the pH level during the FP-biotin labelling reaction and the FP-biotin concentration are the key parameters for sample preparation and merit further optimization.

**Table 1.** Plackett-Burman design with six factors at two levels and twenty runs for the optimization of ABPP sample preparation.

| Run | pH | [FP-biotin] / μM | Labeling time / h | Bead volume / μL | [TCEP] / mM | [IA] / mM |
| --- | --- | --- | --- | --- | --- | --- |
| 1 | 5 | 10 | 2 | 20 | 0 | 20 |
| 2 | 8 | 10 | 2 | 20 | 10 | 0 |
| 3 | 8 | 1 | 2 | 40 | 0 | 0 |
| 4 | 5 | 1 | 2 | 40 | 10 | 20 |
| 5 | 5 | 1 | 2 | 40 | 0 | 20 |
| 6 | 8 | 10 | 0.5 | 20 | 10 | 20 |
| 7 | 8 | 10 | 0.5 | 40 | 0 | 20 |
| 8 | 5 | 1 | 0.5 | 20 | 0 | 0 |
| 9 | 8 | 1 | 0.5 | 20 | 0 | 20 |
| 10 | 8 | 10 | 2 | 40 | 0 | 20 |
| 11 | 5 | 10 | 2 | 20 | 10 | 20 |
| 12 | 5 | 1 | 0.5 | 40 | 10 | 0 |
| 13 | 5 | 10 | 0.5 | 20 | 0 | 0 |
| 14 | 5 | 10 | 2 | 40 | 10 | 0 |
| 15 | 8 | 1 | 2 | 20 | 0 | 0 |
| 16 | 5 | 1 | 0.5 | 20 | 10 | 20 |
| 17 | 8 | 1 | 2 | 20 | 10 | 0 |
| 18 | 8 | 10 | 0.5 | 40 | 10 | 0 |
| 19 | 5 | 10 | 0.5 | 40 | 0 | 0 |
| 20 | 8 | 1 | 0.5 | 40 | 10 | 20 |

**Figure 2.**
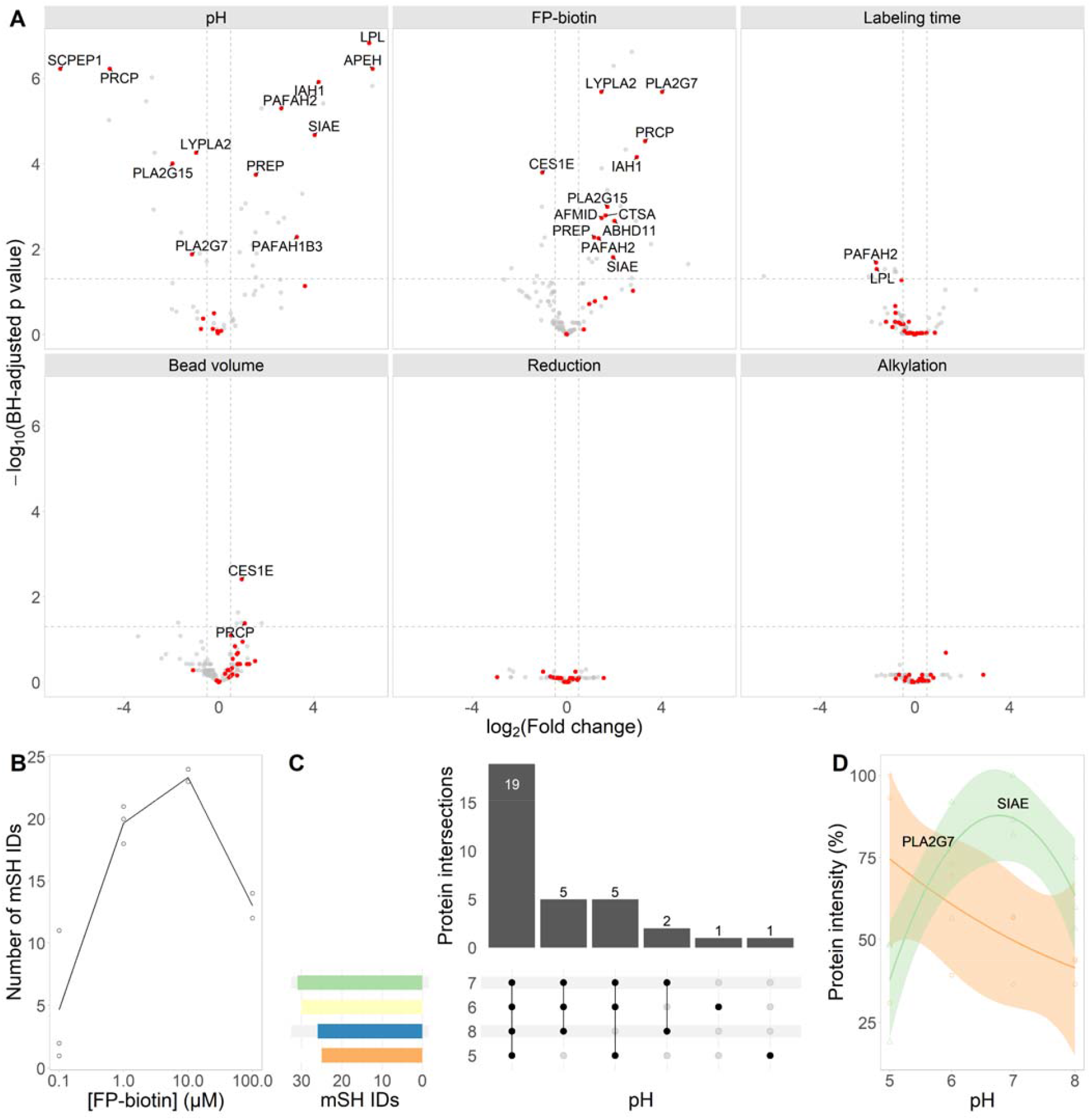
Optimization of ABPP sample preparation. (A) Six parameters were screened according to **Table 1**. Volcano plots indicate how individual mSHs were impacted by the parameter change. Horizontal dashed lines represent adjusted p value < 0.05. Vertical dashed lines represent |log_2_(fold change)| > 0.5. (B) Number of mSH identifications at different FP-biotin concentrations. (C) Number and overlap of mSH IDs at different pH levels. (D) Relative quantification of representative PSDEs at different pH levels.. mAb1 HCCF. (B–D) n = 3.

Accordingly, we optimized FP-biotin concentration and pH level. First, we varied FP-biotin concentration from 0.1 μM to 100 μM in mAb1 HCCF at pH 8. As expected, increasing concentrations resulted in more mSH identifications and higher protein intensity up to 10 μM (**Figure 2B**). However, further increase in the FP-biotin concentration reduced protein identification and intensity. Perhaps, some of the excess FP-biotin or its degradant passed through desalting spin column and competed for streptavidin beads. Second, we varied the pH level from 5 to 8 while fixing the FP-biotin concentration at 10 μM. FP-biotin labelling at pH 7 yielded the highest number of mSH identifications, which was closely followed by pH 6 by one identification (**Figure 2C**). Protein intensity for the majority of known PSDEs, namely LPL, LIPA, PLA2G15 and SIAE, peaked between pH 6 and pH 7, which is slightly higher than the pH levels of most biopharmaceutical formulations (**Figure 2D**). On the other hand, somewhat lower pH was preferable for PLA2G7 while higher pH levels were preferable for several other PSDEs such as CES1 homologs and IAH1. Indeed, a recent study reported optimal pH levels for hydrolytic activity of several purified recombinant PSDEs and confirmed our observations.^31^ These results suggest that performing the FP-biotin labeling step at pH 6 would increase the chance of identifying active and relevant PSDEs in biopharmaceutical formulations.

The combined use of short gradients on Evosep One LC and the fast-scanning PASEF methodology on timsTOF Pro 2 allows for increased sample throughput without compromising ID coverage or data quality.^6,32^ For example, Jones et al. reported a considerable increase in sample throughput in the ABPP of deubiquitylating enzymes in cell and tissue samples using Evosep One LC coupled with timsTOF Pro compared to a conventional nano-flow LC setup with an Orbitrap MS.^33^ Thus, we evaluated several Evosep methods of varying throughput from 200 SPD to 30 SPD in combination with one of the default DDA PASEF methods (“DDA PASEF-short_gradient_0.5sec_cycletime.m”) on timsTOF Pro 2 and benchmarked against the conventional nano-flow LC-MS setup consisting of EASY-nLC 1200 and Orbitrap Q Exactive HF-X (**Figure 3**). The number of mSH identifications increased as the gradient became longer and surpassed the level of identification coverage achievable by the conventional setup despite using a gradient that is less than half in length (44 min vs 110 min), representing nearly four-fold improvement in throughput (30 SPD vs ∼ 8 SPD). In addition, Evotip can load, concentrate, and clean up a relatively large volume (e.g., 20–200 μL) of sample and no longer requires offline clean up, concentration, or reconstitution of samples, which simplifies the protocol and minimizes sample loss. These results demonstrate the advantage of combining a short-gradient LC and the PASEF methodology in ABPP of biologics process intermediates.

**Figure 3.**
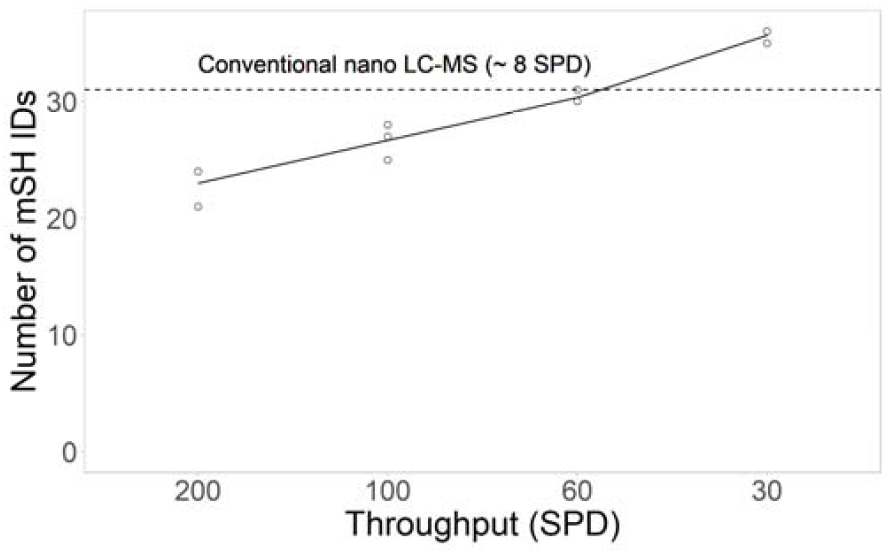
Optimization of Evosep One LC separation method. Conventional nano LC-MS refers to the EASY-nLC 1200– Orbitrap Q Exactive HF-X system. mAb1 HCCF. n = 3.

Previous studies have demonstrated that increasing the TIMS accumulation and ramp times allows fragmenting more precursor ions per PASEF scan and improve sensitivity.^32^ For example, Mun et al. reported the benefit of longer TIMS accumulation and ramp times (180 ms) in single-cell proteomics applications.^34^ On the other hand, the number of PASEF scans should be varied to control for cycle time and sufficient number of data points per peak for quantification.^35^ Thus, we prepared ABPP samples from mAb1 PAP and analyzed these samples under various settings for the TIMS accumulation and ramp times and the number of PASEF scans. Specifically, we applied an eleven-run central composite design^30,36^ centered on the standard DDA PASEF settings and systematically varied the TIMS accumulation and ramp times from the standard values 100 ms to values between 30 ms and 170 ms as well as the number of PASEF scans from the standard value 10 to values between 3 to 17, while operating the instrument near 100% duty cycle (**Table 2**). Indeed, increasing TIMS accumulation and ramp times significantly improved the number of peptide identifications (**Figure 4A**) and representative protein intensity (**Figure 4B**), whereas the number of PASEF scans did not significantly affect either of these measures. In an experiment with somewhat small amount (50 ng) of HeLa protein digest, we observed a similar trend but more pronounced effect of the number of PASEF scans than in the experiment with the ABPP sample, presumably because of increased sample complexity (**Supporting Information Figure S1**). On the other hand, the median number of data points per precursor peaks strongly depended on the TIMS accumulation and ramp times as well as the number of PASEF scans, reflecting the cycle time (**Figure 4C**). These results suggest that DDA PASEF methods for the analysis of biologics ABPP samples should have longer accumulation and ramp times (e.g., 200 ms) as well as smaller number of PASEF scans (e.g., 5 scans) than the standard DDA PASEF method to increase sensitivity while maintaining cycle time (e.g., 1.24 s), hence quantitative accuracy and precision. Analogously, increasing injection time and automated gain control target values on Orbitrap-based instruments would improve sensitivity.^37^

**Table 2.** Central composite design for the optimization of TIMS parameter settings for the DDA PASEF analysis of ABPP samples.

| Run | Accumulation time (ms) | Ramp time (ms) | Number of PASEF scans | Overlap | Cycle time (s) |
| --- | --- | --- | --- | --- | --- |
| 1 | 50 | 50 | 5 | 1 | 0.34 |
| 2 | 150 | 150 | 5 | 1 | 0.94 |
| 3 | 50 | 50 | 15 | 4 | 0.90 |
| 4 | 150 | 150 | 15 | 4 | 2.50 |
| 5 | 100 | 100 | 10 | 4 | 1.17 |
| 6 | 100 | 100 | 10 | 4 | 1.17 |
| 7 | 30 | 30 | 10 | 4 | 0.40 |
| 8 | 170 | 170 | 10 | 4 | 1.94 |
| 9 | 100 | 100 | 3 | 1 | 0.42 |
| 10 | 100 | 100 | 17 | 4 | 1.91 |
| 11 | 100 | 100 | 10 | 4 | 1.17 |

**Figure 4.**
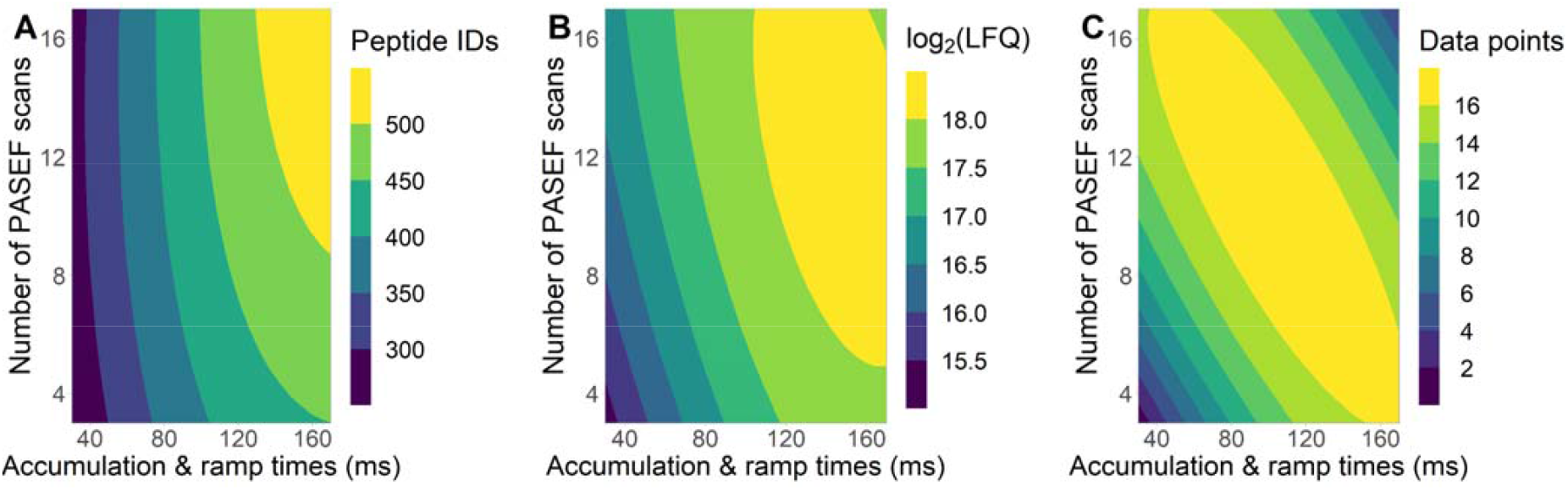
Optimization of TIMS parameter settings for DDA PASEF analysis of ABPP samples. TIMS accumulation and ramp times and number of PASEF scans were varied according to **Table 2**. Response surface models of second order were fitted to (A) number of peptide identifications, (B) label-free quantification of PLA2G7, or (C) median number of data points per peaks (“Number of scans” column of “evidence.txt” file generated by MaxQuant).

Having optimized sample preparation, LC method, and DDA PASEF method, we next evaluated the detection limit of this new workflow. To this end, we spiked mAb1 DS, which lacks measurable PS degradation activity (data not shown), with 1, 10, 100, or 1000 ppb of the purified recombinant PSDEs PLA2G7 and SIAE proteins. Gratifyingly, PLA2G7 was identified between 10–1000 ppb and SIAE was identified between 1–1000 ppb (**Figure 5A**), which was strongly supported by confident identifications of multiple (four for PLA2G7 at 10 ppb and five for SIAE at 1 ppb) unique peptides with highquality spectra in all replicates (**Figure 5B**). This result updates the lowest limit of detection in literature (80 ppb) by eight-fold while using a commercially available chemical reporter.

**Figure 5.**
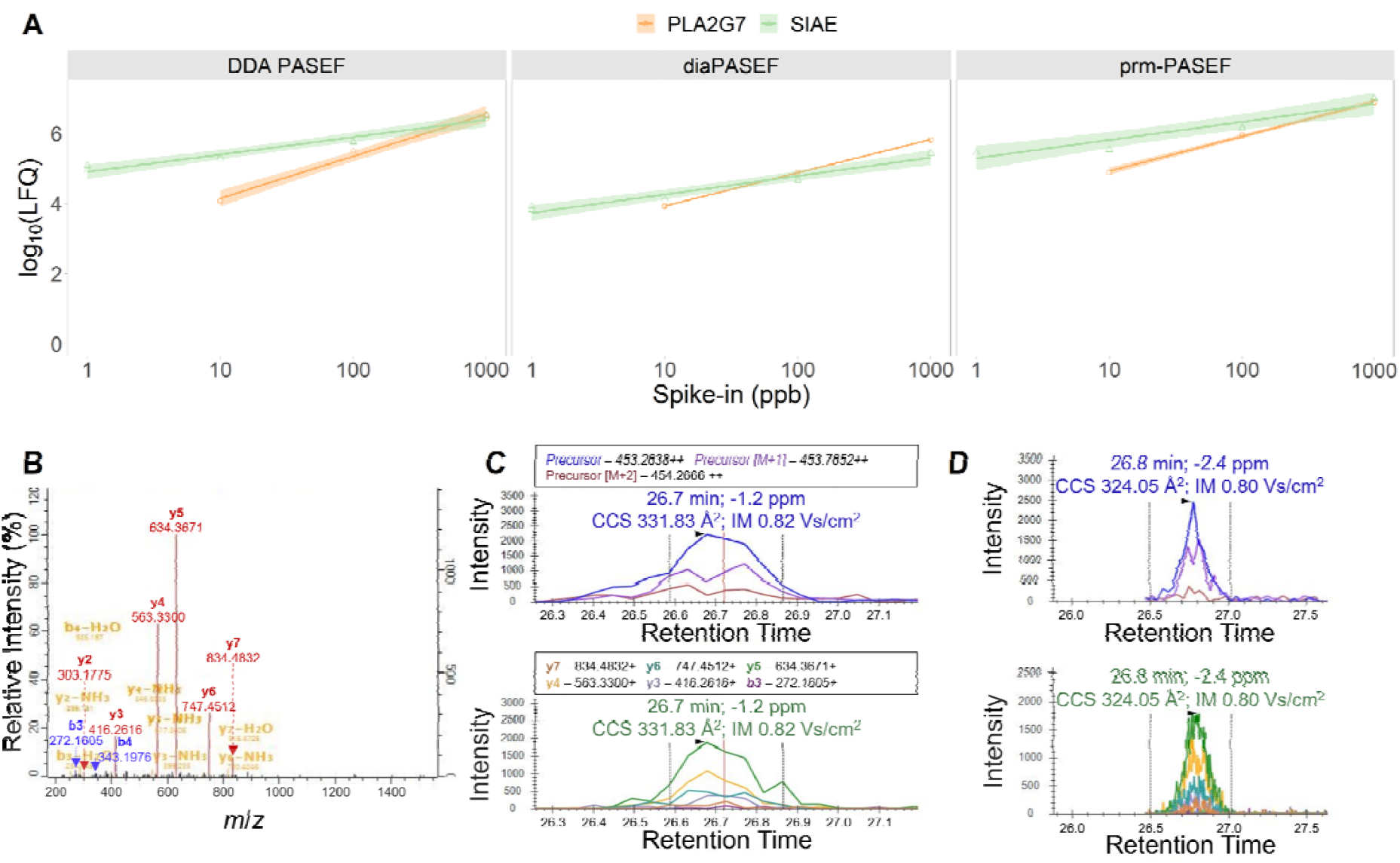
Limit of detection of new ABPP workflow. Purified recombinant PLA2G7 and SIAE were spiked in mAb1 DS at different concentrations of 1, 10, 100, or 1000 ppb. (A) Relative quantification of PLA2G7 and SIAE with three different MS data acquisition modes: DDA PASEF, diaPASEF, or prm-PASEF. Representative MS/MS spectrum or extracted ion chromatograms for the PLA2G7-derived peptide ASLAFLQR detected at the spike-in concentration of 10 ppb with (B) DDA PASEF, (C) diaPASEF, or (D) prm-PASEF. n = 2.

To assess the potential for further enhancing sensitivity by employing alternative data acquisition modes that may be less limited by intra-scan dynamic range, we evaluated the diaPASEF^38^ and prm-PASEF^39^ acquisition modes. We performed a diaPASEF analysis with a general-purpose acquisition scheme^6^ and processed the data with DIA-NN using a spectral library generated by refining an *in-silico* predicted spectra with experimental spectra from the diaPASEF analysis of the mAb1 HCCF ABPP samples, after evaluating multiple data processing strategies (**Supporting Information Figure S2**). For example, the diaPASEF analysis of mAb1 PAP-derived ABPP samples combined with our best data processing strategy led to the identification of 20 additional mSHs that were missed by the DDA PASEF analysis of the same ABPP sample. The prm-PASEF method targeted all precursors that were identified for PLA2G7 and SIAE in the DDA PASEF analysis of the mAb1 HCCF samples (**Supporting Information Table S1**). Both diaPASEF and prm-PASEF identified PLA2G7 down to 10 ppb and SIAE down to 1 ppb with multiple unique peptides supported by high-quality spectra (**Figures 5C & 5D**) and trended to be more precise than DDA PASEF by approximately five percentage points in terms of coefficient of variance. Both methods could be further refined to boost sensitivity and expand the coverage. For example, focusing the diaPASEF method on a smaller mass range^40^ would allow the use of narrower windows and/or longer TIMS accumulation and ramp times. Alternatively, advanced varieties of PASEF-based DIA methods such as Slice-PASEF^41^, which reportedly outperforms diaPASEF for low sample amounts, should be explored. As for the prm-PASEF method, collision energy optimization could potentially significantly improve sensitivity.^42^ In addition, expanding the targets beyond PLA2G7 and SIAE to include other PSDEs is desirable.

Finally, we leveraged the optimized workflow in a root-cause investigation of PS degradation during the mAb2 process development. mAb2 UFP was incubated with PS-80 for two months and residual PS-80 was monitored with an HPLC-CAD assay according to our standard protocol (**Supporting Information Figure S3**). The material exhibited considerable PS-80 degradation, especially at the elevated temperatures of 25 °C and 40 °C, prompting an investigation. To identify potential enzymatic cause of the PS degradation, we performed an ABPP experiment using the new workflow. For comparison, we also conducted an additional ABPP experiment using our original workflow^13^, which requires 20 mg of input per sample, performs FP-biotin labeling at pH 8.0, and acquires data on the conventional nanoflow LC–MS setup with a 110-min gradient (**Table 3**). With the original workflow, we were only able to identify SIAE as an active enzyme with one unique peptide. Unfortunately, SIAE is unlikely to be the cause of observed PS-80 degradation because SIAE hydrolyses PS-20 but not PS-80.^43–45^ In contrast, the new workflow enabled us to identify IAH1 (**Figure 6A**) and SIAE (**Figure 6B**) with multiple unique peptides supported by high-quality spectra. From the standard curb in **Figure 5A**, the amount of SIAE in this material was estimated to be 0.03 ppb (95% confidence interval: 0–0.07 ppb). While IAH1 hydrolyses PS-20, whether it also hydrolyses PS-80 remains unknown and warrants further investigation.^31,43^ Nevertheless, these results clearly demonstrate that the new workflow outperformed the original method and will advance our understanding the root causes of PS degradation.

**Table 3.**
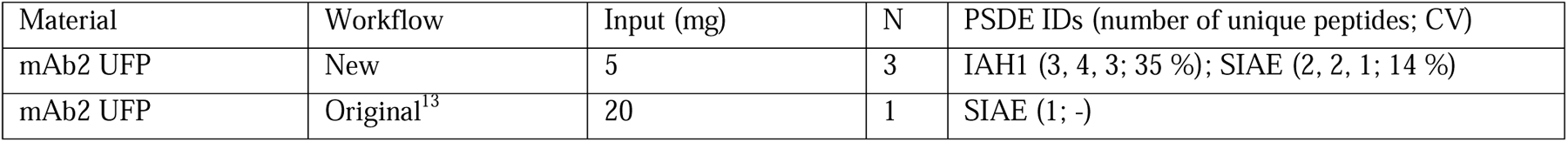
Summary of identified PSDEs by original and new ABPP workflow.

**Figure 6.**
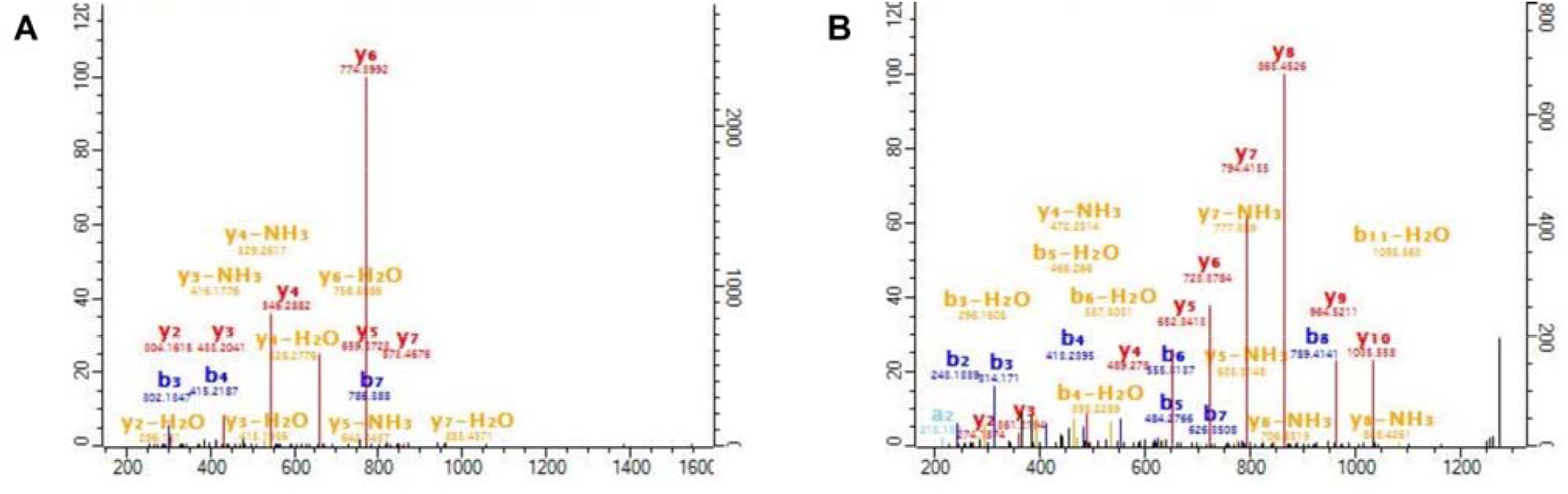
Identification of potential PSDEs in mAb2 UFP. Representative spectra for (A) IAH1-derived peptide SVDILEER and (B) SIAE-derived peptide ELAVAAAYQSVR. n = 3.

## Conclusions

We systematically optimized the FP-biotin–based ABPP workflow for challenging biotherapeutics samples. We comprehensively evaluated every step of the workflow from sample preparation to LC separation and MS data acquisition methods and achieved the identification of as low as 10 ppb of several PSDEs in DS. The new workflow represents the most sensitive analytical tool for PSDE identification and will play critical roles in understanding the cause of PS degradation. Its enhanced sensitivity as well as reduced sample requirement and instrument time has already proved to be advantageous over the original method^13^ in a root-cause investigation. The identification of these trace-level enzymes will guide targeted process approaches to control PS degradation, such as downstream purification, upstream cell culture optimization, and potential genetic knockout during cell line development. Our future work will optimize and extend the targeted proteomics approach for full coverage of known PSDEs. In summary, the new workflow should advance the understanding of biopharmaceutical process and facilitate process and product development.

## ASSOCIATED CONTENT

### Supporting Information

The Supporting Information is available free of charge on the ACS Publications website.

Additional experimental details and results: Optimization of TIMS parameter settings for the DDA PASEF analysis of 50ng HeLa protein digest (**Supporting Information Figure S1**); Evaluation of data processing strategies for the diaPASEF analysis of ABPP samples (**Supporting Information Figure S2**); Polysorbate-80 degradation assay in mAb2 UFP (**Supporting Information Figure S3**); Scheduling list for the prm-PASEF analysis of PLA2G7 and SIAE in mAb1 DS (**Supporting Information Table S1**) (Microsoft Word Document).

## AUTHOR INFORMATION

### Author Contributions

The manuscript was written through contributions of all authors. All authors have given approval to the final version of the manuscript.

## ACKNOWLEDGMENT

TT thanks Merck & Co., Inc., Rahway, NJ, USA Postdoctoral Program. The authors thank Kaniz Fatema for her help with executing experiments. The authors thank Douglas D. Richardson and Hillary A. Schuessler for guidance and support. The authors thank our internal Polysorbate Degradation and Lipase Control Strategy working group for discussions.

## For Table of Contents Only

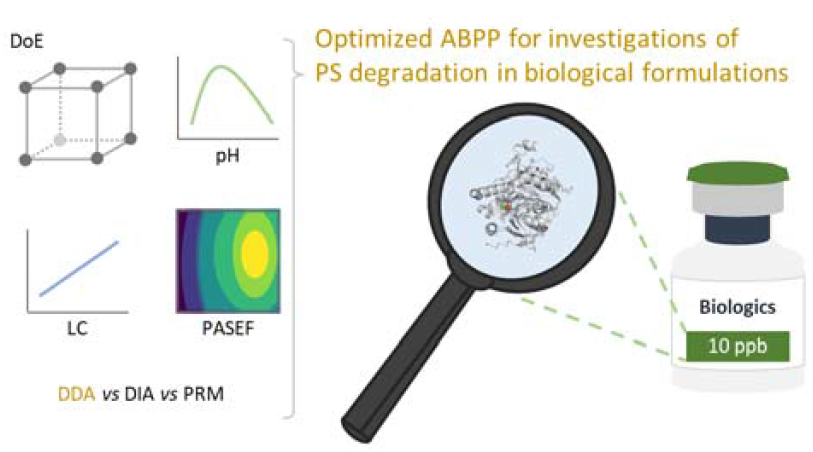

