## Supporting Information for "Systematic Optimization of Activity-Based Protein Profiling for Identification of Polysorbate-Degradative Enzymes in Biotherapeutic Drug Substance Down to 10 ppb"

**Contents**

Figure S1………………………………………………………………………………………………S2

Figure S2………………………………………………………………………………………………S3

Figure S3………………………………………………………………………………………………S4

Table S1….……………………………………………………………………………………………S5

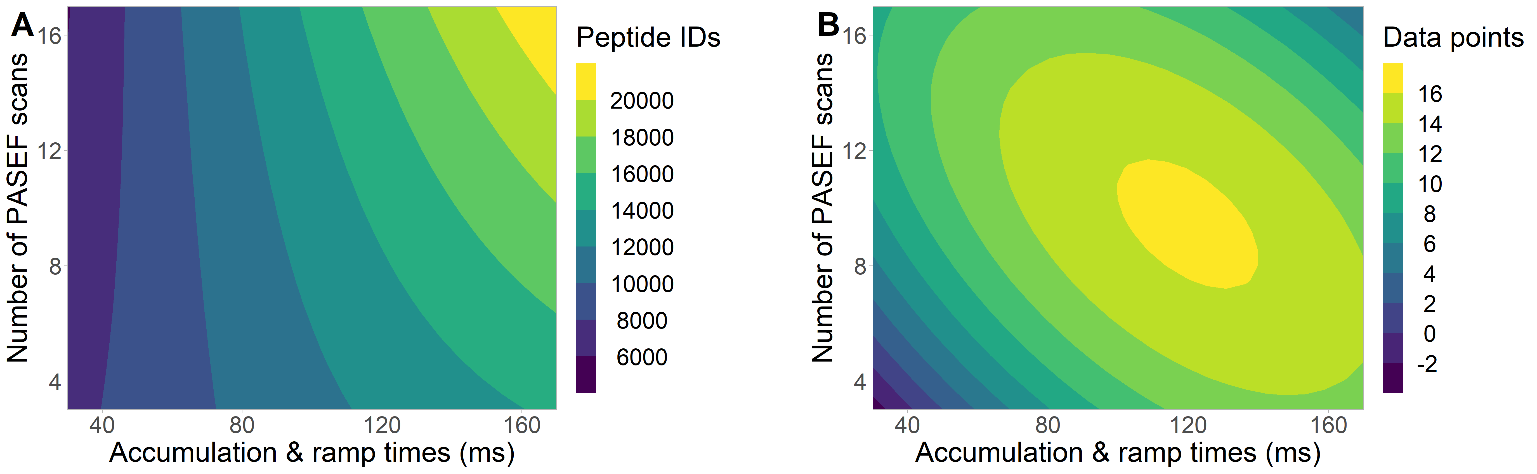

**Figure S1.** Optimization of TIMS parameter settings for the DDA PASEF analysis of 50ng HeLa protein digest. TIMS accumulation and ramp times and number of PASEF scans were varied according to **Table 2**. Response surface models of second order were fitted to (A) number of peptide identifications, or (B) median number of data points per peaks (“Number of scans” column of “evidence.txt” file generated by MaxQuant).

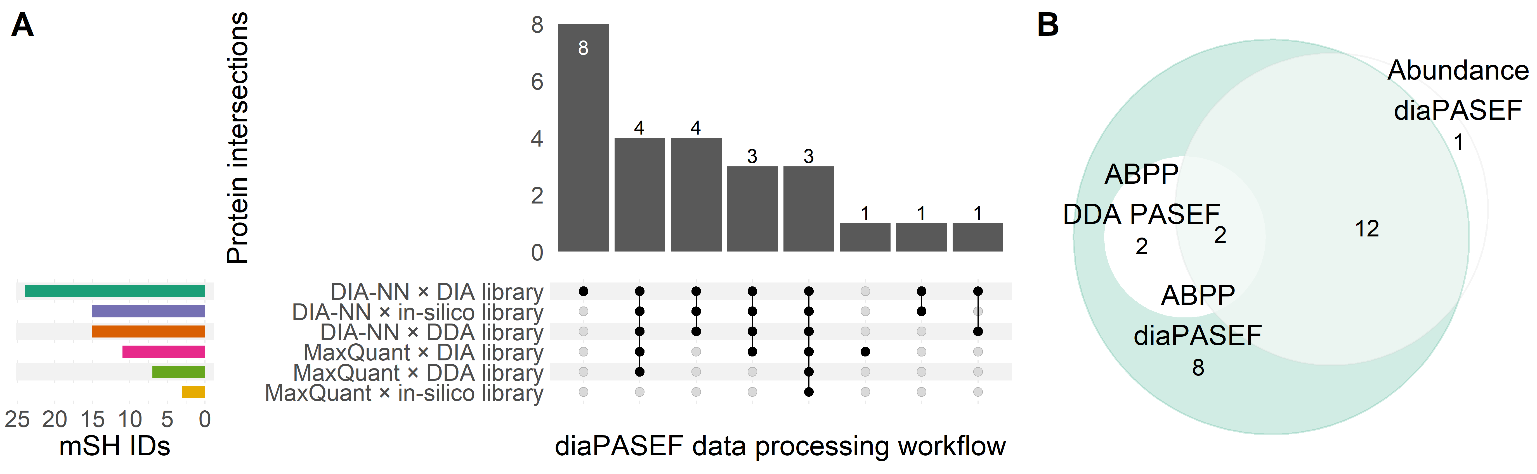

**Figure S2.** Evaluation of data processing strategies for the diaPASEF analysis of ABPP samples. (A) Comparison of mSH IDs among various data processing strategies. DIA-NN or MaxQuant was used in combination with different spectral libraries. DDA library: DDA PASEF analysis of a mAb1 HCCF sample with MaxQuant; in-silico library: *in-silico* prediction on MaxQuant or DIA-NN; DIA library: refinement of the *in-silico* library with diaPASEF analysis of the mAb1 HCCF sample on MaxQuant or DIA-NN. (B) Comparison of mSH IDs among the DDA PASEF analysis of mAb1 PAP–derived ABPP samples (ABPP DDA PASEF), the diaPASEF analysis of mAb1 PAP–derived ABPP samples (ABPP diaPASEF), and the diaPASEF analysis of mAb1 PAP native digestion sample (Abundance diaPASEF). The mSH uniquely identified by Abundance diaPASEF is SEC23IP, which belongs to the PA-PL1 family but seems to lack phospholipase activity according to UniProt.

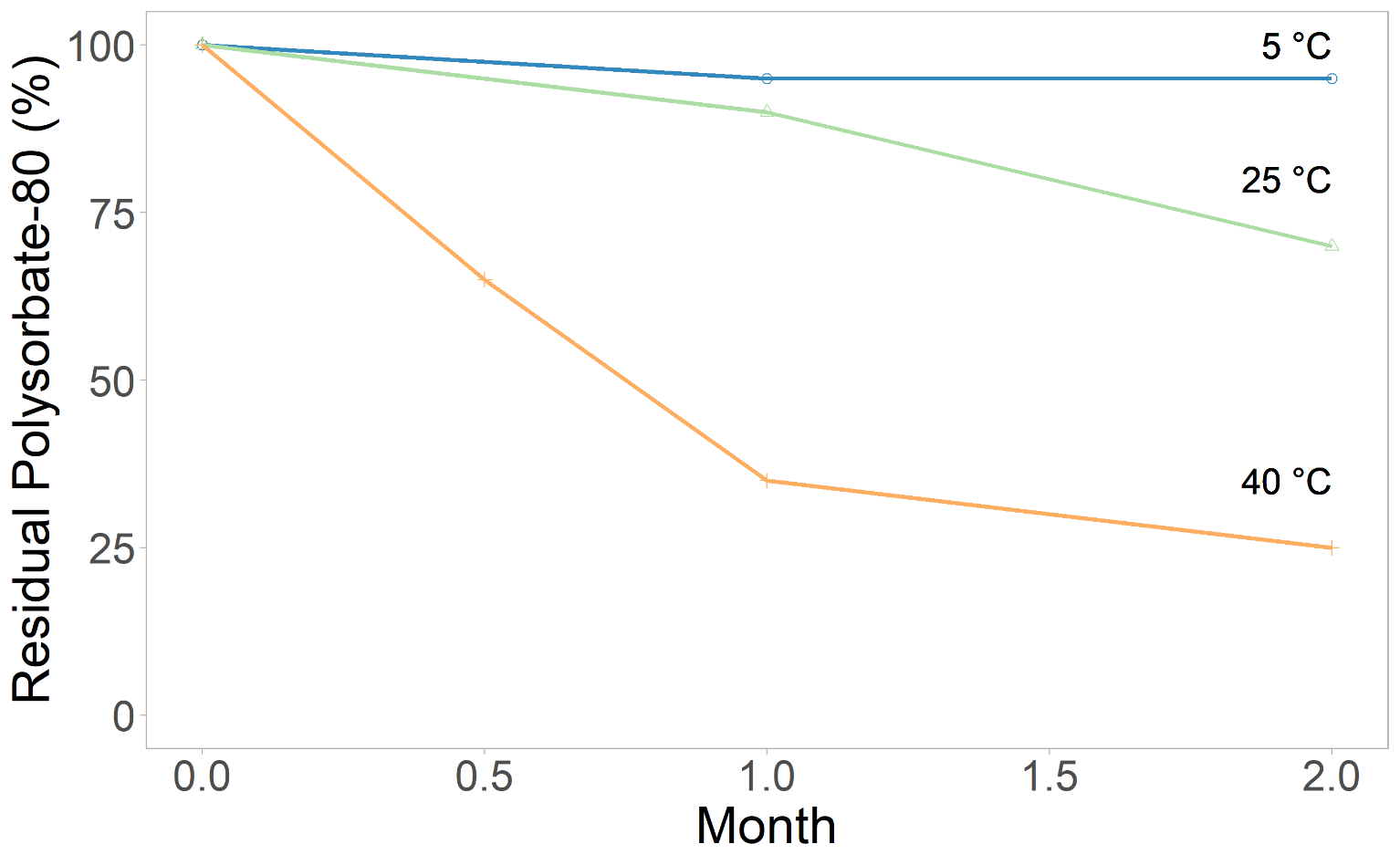

**Figure S3.** Polysorbate-80 degradation assay in mAb2 UFP.

**Table S1.** Scheduling list for the prm-PASEF analysis of PLA2G7 and SIAE in mAb1 DS.

| Charge | Mass [m/z] | Isolation Width [m/z] | RT [s] | RT Range [s] | Start IM [1/K0] | End IM [1/K0] | CE [eV] | External ID | Description |
| --- | --- | --- | --- | --- | --- | --- | --- | --- | --- |
| 2 | 453.2638 | 3 | 1647.69 | 120 | 0.758047 | 0.858047 | 27.68523 | ASLAFLQR | PLA2G7 |
| 2 | 1102.008 | 3 | 2424.54 | 120 | 1.207133 | 1.307133 | 45.43956 | CGVSLDPWMYPVSEELYSK | PLA2G7 |
| 2 | 719.3359 | 3 | 2071.8 | 120 | 0.95269 | 1.05269 | 35.5198 | DFNQWDSLVEGK | PLA2G7 |
| 2 | 911.9101 | 3 | 2054.76 | 120 | 1.083982 | 1.183982 | 40.81931 | DGSASATYYFEDQVAAK | PLA2G7 |
| 2 | 913.4593 | 3 | 2100.36 | 120 | 1.194848 | 1.294848 | 44.8255 | ECSQALSAILDIGHGSPK | PLA2G7 |
| 2 | 642.8272 | 3 | 2110.92 | 120 | 0.915391 | 1.015391 | 33.95891 | ENILGSYFDVK | PLA2G7 |
| 2 | 495.7502 | 3 | 1881.6 | 120 | 0.803116 | 0.903116 | 29.33168 | EYFFGLSK | PLA2G7 |
| 2 | 716.3699 | 3 | 1963.8 | 120 | 0.945235 | 1.045235 | 35.28465 | GEIDSGVAIDLTNK | PLA2G7 |
| 3 | 585.6214 | 3 | 1858.92 | 120 | 0.813119 | 0.913119 | 29.3065 | GSVHQNFADFTFVTGK | PLA2G7 |
| 2 | 981.5278 | 3 | 2337.3 | 120 | 1.172719 | 1.272719 | 43.33243 | IPQPLFFINSAQFQSPK | PLA2G7 |
| 2 | 496.7633 | 3 | 676.29 | 120 | 0.783097 | 0.883097 | 28.50414 | IQAVMSTAR | PLA2G7 |
| 2 | 543.3031 | 3 | 1593.48 | 120 | 0.813119 | 0.913119 | 30.47861 | LDTVWIPNK | PLA2G7 |
| 2 | 619.296 | 3 | 998.64 | 120 | 0.887998 | 0.987998 | 32.51733 | LYYPAQDPDR | PLA2G7 |
| 3 | 916.1263 | 3 | 1937.58 | 120 | 0.977522 | 1.077522 | 36.44307 | NLIPGSPSDVVNLSPTLQSSPGSHTQN | PLA2G7 |
| 2 | 559.2722 | 3 | 705.12 | 120 | 0.823117 | 0.923117 | 30.29703 | QLEGAIDEDK | PLA2G7 |
| 2 | 431.7139 | 3 | 832.05 | 120 | 0.73046 | 0.83046 | 26.85491 | SSAWFHK | PLA2G7 |
| 2 | 788.9354 | 3 | 2139.78 | 120 | 1.060506 | 1.160506 | 39.61683 | YPLVIFSHGLGAFR | PLA2G7 |
| 2 | 639.3461 | 3 | 1550.64 | 120 | 0.917879 | 1.017879 | 34.10479 | ELAVAAAYQSVR | SIAE |
| 3 | 989.4954 | 3 | 2473.08 | 120 | 1.172719 | 1.272719 | 44.06394 | EPSGAVIWGFGTPGATVTVTLCQGQNTFK | SIAE |
| 2 | 789.3871 | 3 | 1952.1 | 120 | 1.032054 | 1.132054 | 38.30941 | FGSYIDNYMVLQK | SIAE |
| 2 | 373.2138 | 3 | 562.824 | 120 | 0.7 | 0.790273 | 25.05955 | GTVVDVR | SIAE |
| 2 | 890.4185 | 3 | 1811.04 | 120 | 1.059269 | 1.159269 | 39.62871 | GVIWYQGESNEDLNR | SIAE |
| 2 | 469.7745 | 3 | 815.7 | 120 | 0.793109 | 0.893109 | 29.15202 | IELLAHSR | SIAE |
| 2 | 1018.507 | 3 | 2477.04 | 120 | 1.167799 | 1.267799 | 43.74752 | IFSASLVQAEEELEDLDK | SIAE |
| 2 | 540.8004 | 3 | 1402.47 | 120 | 0.839355 | 0.939355 | 31.16785 | LLSLTYDQK | SIAE |
| 2 | 626.3419 | 3 | 2167.44 | 120 | 0.875536 | 0.975536 | 32.4771 | MTVLQIFNASK | SIAE |
| 2 | 542.8111 | 3 | 1517.91 | 120 | 0.835609 | 0.935609 | 31.12595 | NLAFQGPLPK | SIAE |
| 2 | 524.7602 | 3 | 1955.58 | 120 | 0.800615 | 0.900615 | 29.70495 | PGGPYEVMAK | SIAE |
| 2 | 625.3486 | 3 | 1706.64 | 120 | 0.902944 | 1.002944 | 33.47195 | QTFGITNVTLR | SIAE |
| 2 | 524.2463 | 3 | 248.382 | 120 | 0.76807 | 0.86807 | 28.22475 | QTFHDGSQK | SIAE |
| 2 | 582.2591 | 3 | 439.179 | 120 | 0.808118 | 0.908118 | 29.73994 | SCGVPNTSNER | SIAE |
| 2 | 431.7245 | 3 | 1362.54 | 120 | 0.73046 | 0.83046 | 26.78668 | VDLSWSR | SIAE |
| 3 | 553.9475 | 3 | 1830.78 | 120 | 0.778089 | 0.878089 | 28.07673 | WHQTADFGFVPNLK | SIAE |
| 2 | 682.8641 | 3 | 1917.75 | 120 | 0.927831 | 1.027831 | 34.59129 | WLPVSVNSFSTK | SIAE |
| 3 | 1044.517 | 3 | 2514.36 | 120 | 1.046903 | 1.146903 | 38.48102 | YLYDNLQYPIGLVSSSWGGTPIEAWSSK | SIAE |

This table is formatted for the Bruker Hystar software. External ID here refers to the targeted peptide sequence (peptide sequence and charge state, or any unique characters).
